# Spatially Resolved Transcriptomics for Evaluation of Intracranial Vessels in a Rabbit Model: Proof of Concept

**DOI:** 10.1101/2022.02.09.479726

**Authors:** Matthew S. Zabriskie, Daniel L. Cooke, Chuanzhuo Wang, Matthew D. Alexander

## Abstract

**Background:** Better understanding of vessel biology and vascular pathophysiology is needed to improve understanding of cerebrovascular disorders. Tissue from diseased vessels can offer the best data. Rabbit models can be effective for studying intracranial vessels, filling gaps resulting from difficulties acquiring human tissue. Spatially-resolved transcriptomics (SRT) in particular hold promise for studying such models as they build on RNA sequencing methods, augmenting such data with histopathology.

**Methods:** Rabbit brains with intact arteries were flash frozen, cryosectioned, and stained with H&E to confirm adequate inclusion of intracranial vessels before proceeding with tissue optimization and gene expression analysis using the Visium SRT platform. SRT results were analyzed with k-means clustering analysis, and differential gene expression was examined, comparing arteries to veins.

**Results:** Cryosections were successfully mounted on Visium proprietary slides. Quality control thresholds were met. Optimum permeabilization was determined to be 24 minutes for the tissue optimization step. In analysis of SRT data, k-means clustering distinguished vascular tissue from parenchyma. When comparing gene expression traits, the most differentially expressed genes were those found in smooth muscle cells. These genes were more commonly expressed in arteries compared to veins.

**Conclusions:** Intracranial vessels from model rabbits can be processed and analyzed with the Visium SRT platform. Face validity is found in the ability of SRT data to distinguish vessels from parenchymal tissue and differential expression analysis accurately distinguishing arteries from veins. SRT should be considered for future animal model investigations into cerebrovascular diseases.

## Introduction

Cerebrovascular diseases are leading causes of death and long-term disability worldwide, yet there is poor understanding of their pathophysiology and natural history.(1–8) This limited understanding in turn leads to imprecise treatment options and uncertainty about why some treatments work and others do not. The inadequate data on cerebrovascular pathophysiology results from the prohibitive morbidity of obtaining tissues for analysis, which is traditionally considered the gold standard for disease evaluation. To date, management strategies have largely been based on extrapolation of the pathophysiology of diseases in surrogate extracranial vessels. This approach assumes intracranial and extracranial arterial systems share similar physiology and pathology. This assumption is scientifically suspect and should be investigated because these distinct systems are derived from different germ cell layer.(9–17) There is a critical gap in current understanding of the underlying biology of intracranial arteries and the pathophysiologic features of diseases that occur in them. There is a crucial need for better data from intracranial vessels themselves to better understand biological processes in these arteries and the diseases they can develop.

For situations in which tissue cannot be safely sampled for study in humans, model animals can provide opportunities for research. In particular, model animals can allow interrogation of the entire course of diseases. In contrast, in the rare scenarios in which human tissue data can be obtained, these offer a snapshot from only a single point in time. Such human data typically represent advanced, symptomatic disease, while many of these processes develop and progress over years. Additionally, such diseases typically occur in humans with numerous comorbidities that confound analysis, and they frequently exist for years or decades without causing symptoms. In addition to providing opportunities for tissue-level interrogation across all stages of a disease’s natural history, whether symptomatic or not, animal models also allow investigators to control for such confounders to better identify processes specific to the disease of interest.

Rabbits are a particularly valuable model animal for the study of cerebrovascular diseases. They have a long track record of effective analysis for cardiovascular diseases and subsequent application to cerebrovascular diseases with effective modeling of human disease.(18–25) Additionally, their size allows for endovascular access to perform minimally invasive techniques, and absence of a rete mirabile allows for catheterization of the intracranial circulation, which cannot be achieved in most other large model animals.(25–31)

As animal models have traditionally been evaluated with histology and immunohistochemical staining, the rapidly progressing fields of genomics and transcriptomics allow for deeper analysis. RNA sequencing (RNAseq) techniques have evolved from bulk analysis of tissue samples to single-cell techniques that can distinguish gene expression patterns among various cell types. Indeed, members of our group have previously reported endovascular biopsy and single-cell RNAseq results from animal and human samples.(26–28,30,32–34) However, these techniques lack spatial information, so it cannot be known from where certain gene expression findings originate.

While RNAseq techniques yield valuable gene expression data, to date, these methods have been limited by the lack of spatial information.(35,36) Even when evaluating single cell data, the origin of a cell is unknown. Spatially-resolved transcriptomics (SRT) methods overcome this limitation, providing gene expression data for thousands of locations on a pathologic slide. A grid of thousands of spots, measuring less than 10 square microns, overlies a conventional microscopic histopathology image.(37) Referring gene expression to anatomic location increases likelihood of mechanistic insights resulting from investigations of diseases. Early uses of SRT include cortical samples that include small arterial branches, but application for intracranial vessels specifically has not been published.(38) This report describes the methodology for performing SRT on tissue harvested from a rabbit for the express purpose of studying the intracranial vasculature. SRT data can augment findings from single-cell transcriptomic analyses of normal and diseased human vascular tissues.(32–34)

## Methods

### Rabbits

Two apolipoprotein E knockout Oryctolagus cuniculus rabbits from the same litter were selected for euthanasia and spatial transcriptomic analysis.(21) Rabbits have an extensive history of use for studying vascular disorders throughout the body, and the apolipoprotein E knockout model in particular has been used for modeling intracranial disease.(20,21,27,30,34) Both were euthanized at 14 months of age. Rabbit 1 was a male weighing 4.2 kg. Rabbit 2 was a female weighing 4.8 kg.

### Euthanasia

Euthanasia was performed in accordance with a protocol approved by the Institutional Animal Care and Use Committee and previously published.(18,25) General endotracheal anesthesia was induced with intramuscular injection of buprenorphine (0.03 mg/kg) followed later by a ketamine (25-35 mg/kg) and xylazine (3mg/kg) mixture. The fur of the pelvis was trimmed bilaterally. An incision was made to expose and isolate a femoral artery with blunt dissection.(18,19,25) With vessel loops applying gentle traction, the artery was cannulated with a 22-gauge intravenous catheter, through which an 0.018-inch microwire was then advanced into the artery. The catheter was removed over the wire and replaced with a vascular access sheath using Seldinger technique. The sheath introducer and microwire were removed, confirming intra-arterial placement via back bleeding of arterial blood. A midline abdominal incision was then made, followed by blunt dissection to isolate the inferior vena cava (IVC). A steady flow of 0.1M phosphate buffered saline (PBS), pH 7.4, was then infused through the indwelling sheath.(18,19,25) Shortly after initiating perfusion, the IVC was transected to exsanguinate the animal and minimize residual blood in the tissues. After confirming euthanasia, the animal was decapitated, and the brain with intact arteries was harvested.

### Fresh Frozen Block Preparation

The brain surface was then cleaned, dried by blotting gently with delicate task wipes, and sliced into smaller segments for flash freezing. Samples were embedded in optimal cutting temperature (OCT) compound and snap frozen in an isopentane bath submerged in liquid nitrogen. Blocks were then stored at −80°C until slicing was performed at a later date.

### Cryosectioning and Staining

In accordance with the recommended protocol for the Visium SRT platform (10X Genomics, San Francisco, CA), sectioning of blocks was performed with initial cryostat settings of −20°C for the blade and −10°C for the specimen head.(39) 10 µm thick slices were then made, adjusting temperatures as needed to mitigate tearing or curling to obtain the highest quality slices.(39) Representative samples from multiple blocks were stained with hematoxylin and eosin (H&E) to confirm presence of intracranial arteries of interest under light microscopy. Three blocks most suitable for further analysis were then chosen: block 2—coronal section of central right cerebral hemisphere of rabbit 2 with intact internal carotid artery in cross section and middle cerebral artery in longitudinal orientation; block 8—coronal section of central left cerebral hemisphere of rabbit 1 with intact internal carotid artery in cross section; block 11—paramedian sagittal section of left brainstem and cerebellum from rabbit 2 with intact rostral cerebellar artery branches in cross section.(Figure 1) After mounting tissue samples on slides that were deemed adequate for inspection, each block was closed by covering the cut edge in OCT and returning to −80°C without allowing the tissue to thaw.(39)

**Figure 1.**
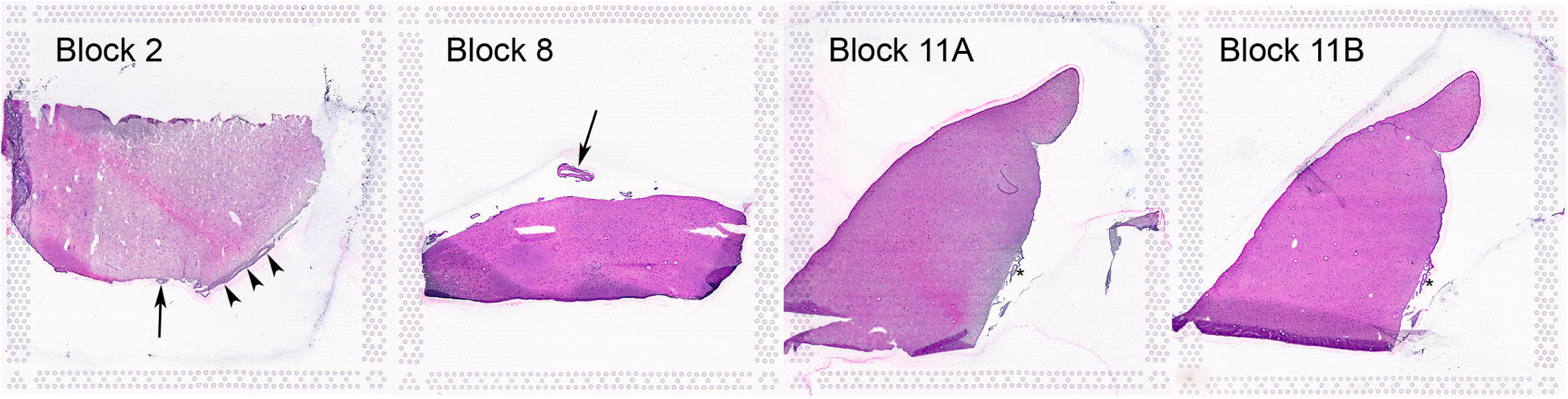
40X light microscopic images of H&E-stained tissue sample on Visium Gene Expression Slide. (Block 2) inferior aspect of coronal plane slice of the central right cerebral hemisphere. (Block 8) inferior aspect of coronal plane slice of the central left cerebral hemisphere. (Blocks 11A/B), sagittal slice of paramedian brainstem and cerebellum. Arrows indicate internal carotid arteries in cross section. Arrowheads indicate middle cerebral artery branch in longitudinal section. Asterisks demonstrate rostral cerebellar arteries in cross section. Note fiducial markers for SRR alignment in the periphery of each block.

### RNA Quality Assessment

In accordance with the Visium protocol, 10 µm thick tissue sections were obtained from blocks 2, 8, and 11 and placed individually in microcentrifuge tubes.(40) The blocks were covered with OCT on the cut edge and returned to −80°C without allowing the block to thaw. RNA extraction was then performed on the sliced samples using the RNeasy Mini protocol for purification of total RNA from animal tissues (Qiagen, Hilden, Germany). Purified RNA was then subjected to RNA integrity number (RIN) calculation using the Agilent RNA 6000 Pico kit (Agilent Technologies, Santa Clara, CA). RINs exceeding the Visium optimization threshold of 7.0 were confirmed for sections from all three blocks of interest.(40)

### Tissue Optimization

SRT analysis performed on the Visium platform involves placement of tissue samples on proprietary microscope slides with capture areas that contain oligonucleotides for mRNA capture. Samples from block 8 were placed on the capture areas and fixed with methanol, stained with H&E, and imaged with light microscopy. The slide-mounted tissues were then permeabilized, and poly-adenylated mRNA was subjected to reverse transcription to generate cDNA.(40) Since different tissues require different permeabilization durations, this was optimized with the proprietary Visium tissue optimization slide.(40) On this slide, the seven tissue samples from block 8 were placed next to an empty control area for eight total sites to evaluate optimum permeabilization time. Six of these sites were permeabilized for different durations (3, 6, 12, 18, 24, or 30 minutes). One capture area was left empty with no tissue, although this capture area contains reference RNA for positive control. Another capture area had a tissue sample placed but was not subjected to permeabilization to serve as a negative control. After the permeabilization step, enzymatic tissue removal was performed, and the remaining fluorescently labeled cDNA underwent fluorescence imaging to assess signal among the various samples. The optimum permeabilization duration was determined by identifying the sample with greatest signal without blurring.

### Gene Expression

Visium gene expression analysis was then performed using the optimal permeabilization duration.(40) The proprietary Visium gene expression slide contains oligonucleotides with spatial barcodes corresponding to a precise physical location on the slide. Each capture area contains ∼5000 spots with unique barcodes. cDNA libraries generated include this geographic barcode information so that gene expression data can be mapped back to the geographic location on the tissue section. Additionally, this can be visually inspected to refer back to the H&E-stained image to localize the transcriptomic data to known anatomic structures. The propriety slide contains four capture areas. For the current analysis, one tissue sample each from blocks 2 and 8, as well as two from block 11, were placed in these separate areas. After slice placement on the slide, permeabilization and reverse transcription were performed following parameters chosen from the tissue optimization step, followed by second strand synthesis and denaturation and cDNA amplification. After passing quality control, a cDNA library was constructed and sequencing performed on the Illumina NovaSeq platform (Illumina, San Diego, CA).

### Vessel Identification with SRT and Differential Expression Analysis

Binary Base Call (BCL) files generated by the Illumina (Illumina, San Diego, CA) platform were converted to FASTQ files using the mkfastq program. Following quality control analysis to confirm data standards were met, data from each of the four capture areas were exported into cloupe files for analysis with Loupe Browser (10X Genomics, San Francisco, CA). Gene expression data were then analyzed to identify the ability of differential expression traits to label different cell types. Before visualizing SRT dots, clearly visible vessels were traced, and then gene expression data for dots within each of the traced regions were isolated. To maximize specificity to vessels, only those on or just adjacent to the brain surface were included. Possible but indeterminate vessel fragments, vessels not clearly distinguishable from parenchyma, and clear vessels located within parenchyma were excluded from tracings in order to maximize specificity of gene expression data from dots representing vessels. Dots were then unhidden, and those within the confines of the traced regions were assigned the label “vessel.” Additionally, further classification was assigned based on whether a dot corresponded to an artery or vein based on histologic appearance. For unsupervised assessment, k-means clustering analysis was performed using n=3 clusters for all tissue and for the vessel-traced regions. Finally, differential expression of dots within vessel tracings was examined, comparing gene expression between arteries and veins.

## Results

Representative images from blocks 2, 8, and 11 are provided in Figure 1. RIN results were 8.70, 7.20, and 7.20 for blocks 2, 8, and 11, respectively. Figure 2 provides electrophoresis and RNA peaks from RNA integrity analysis. After tissue optimization analysis, a 24-minute duration was chosen as the optimum parameter for permeabilization. Figure 3 demonstrates the H&E and fluorescence imaging results from this step.

**Figure 2.**
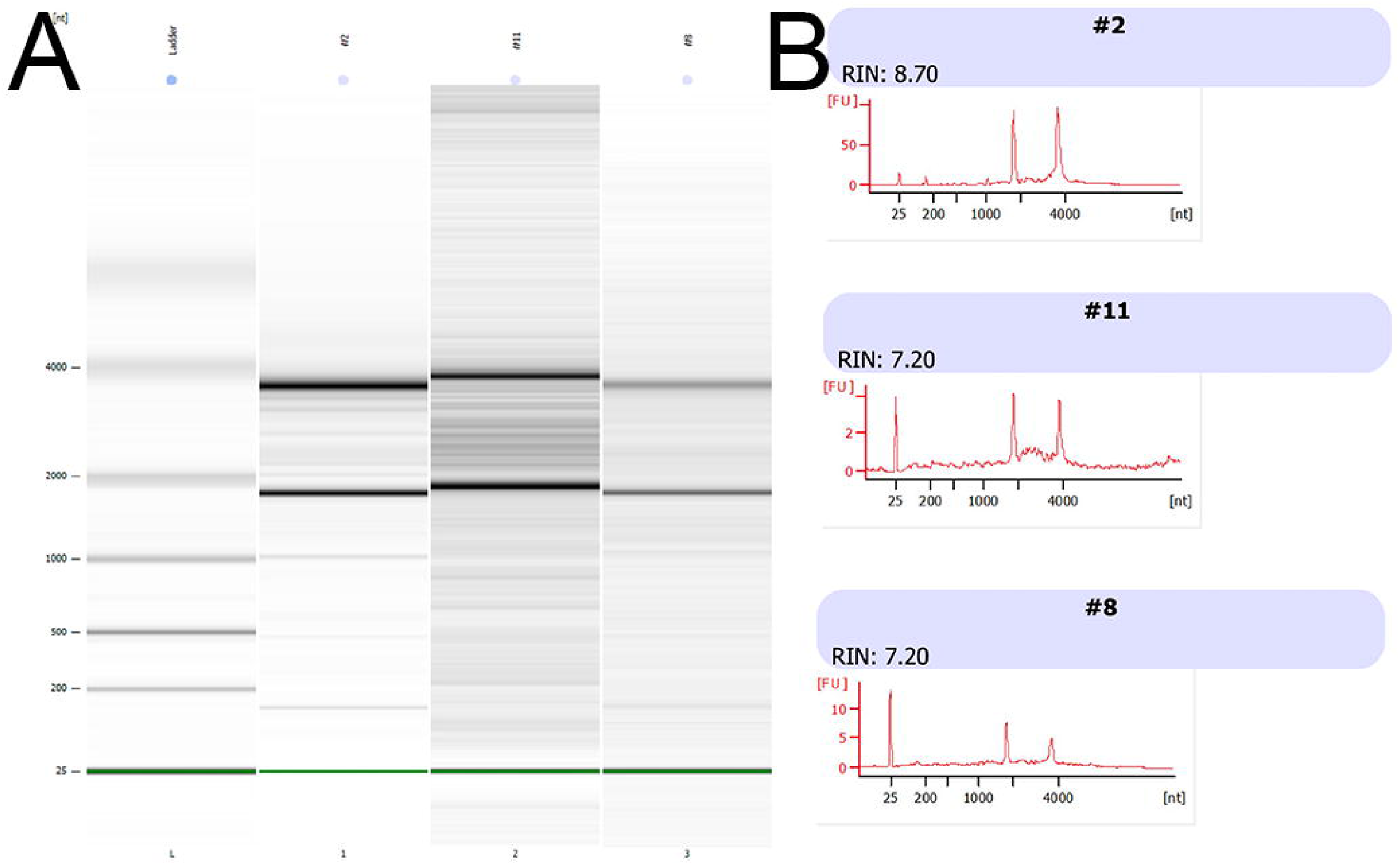
RIN analysis results. (A) Electrophoresis with lanes for control (ladder) and Blocks 2, 11, and 8. (B) RNA peaks from three tissue blocks, with 18S and 28S ribosomal peaks at approximately 1800 nt and 3800 nt, respectively.

**Figure 3.**
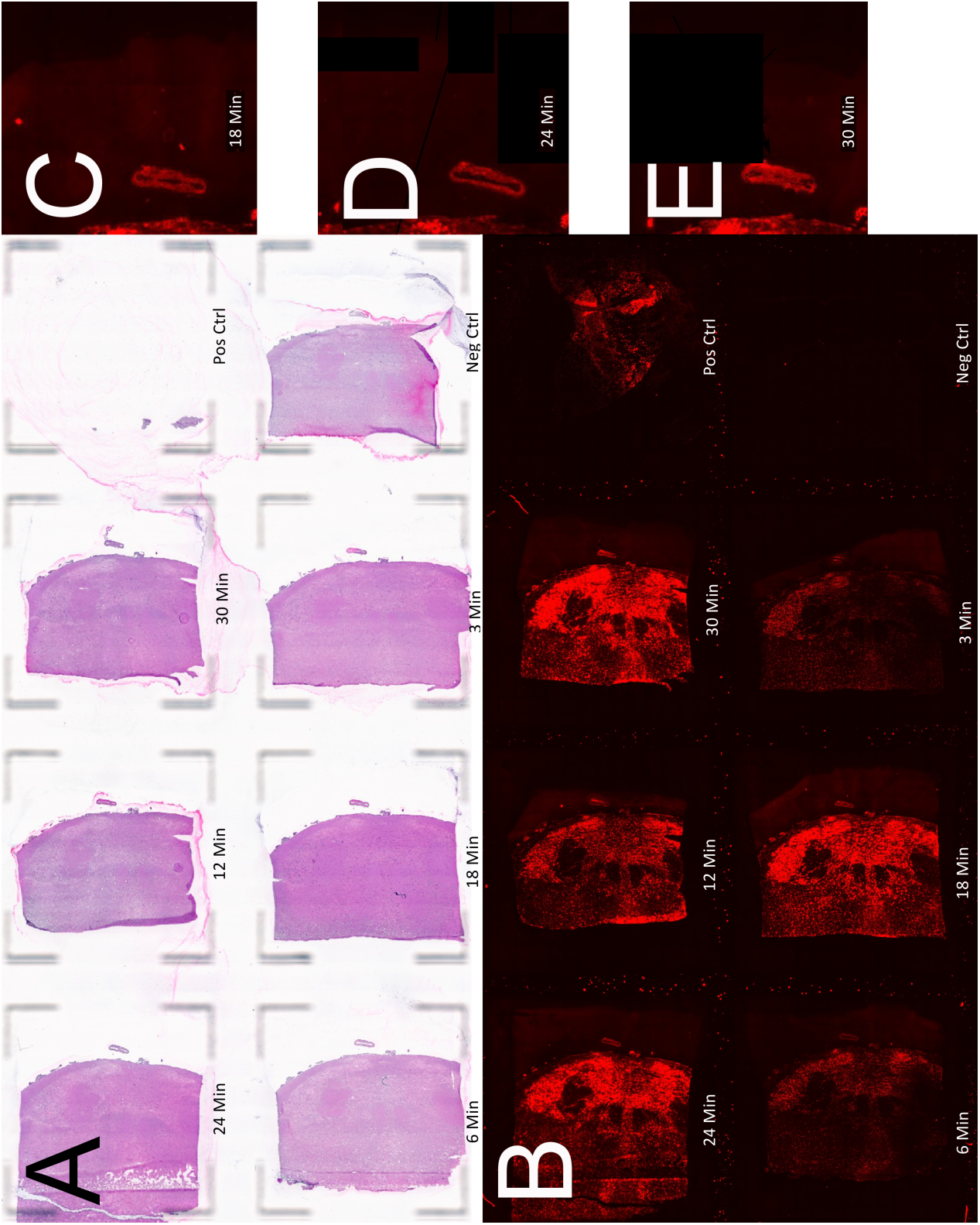
(A) H&E and (B) fluorescence images of the Visium Tissue Optimization slide with slices of Block 8. Permeabilization times are noted below each tissue sample. Magnified images of the left internal carotid artery are provided for 18 minutes (C), 24 minutes (D), and 30 minutes (E). 24 minutes was selected as the optimum permeabilization time for its combination of high signal with minimal blurring.

Figure 4 provides representative images from block 2, showing H&E on the Visium gene expression slide, dots with color-coded K-means cluster assignment, and a fused image. Additionally, Figure 4 provides higher magnification view and color-coded dots based on visual assignment to vessel and artery/vein. Overall, 7039 dots were identified with high quality gene expression data (1940 from Block 2, 1296 from Block 8, 1916 from Block 11A, 1887 from Block 11B). 127 dots were assigned to the cluster corresponding to vessels by K-means clustering (67 from Block 2, 24 from Block 8, 19 from Block 11A, 17 from Block 11B). 69 total dots were included in vessel tracing regions. Among these, 62 (89.9%) were assigned to the K-means cluster corresponding to vessels, while 7 (10.1%) were assigned to other clusters.

**Figure 4.**
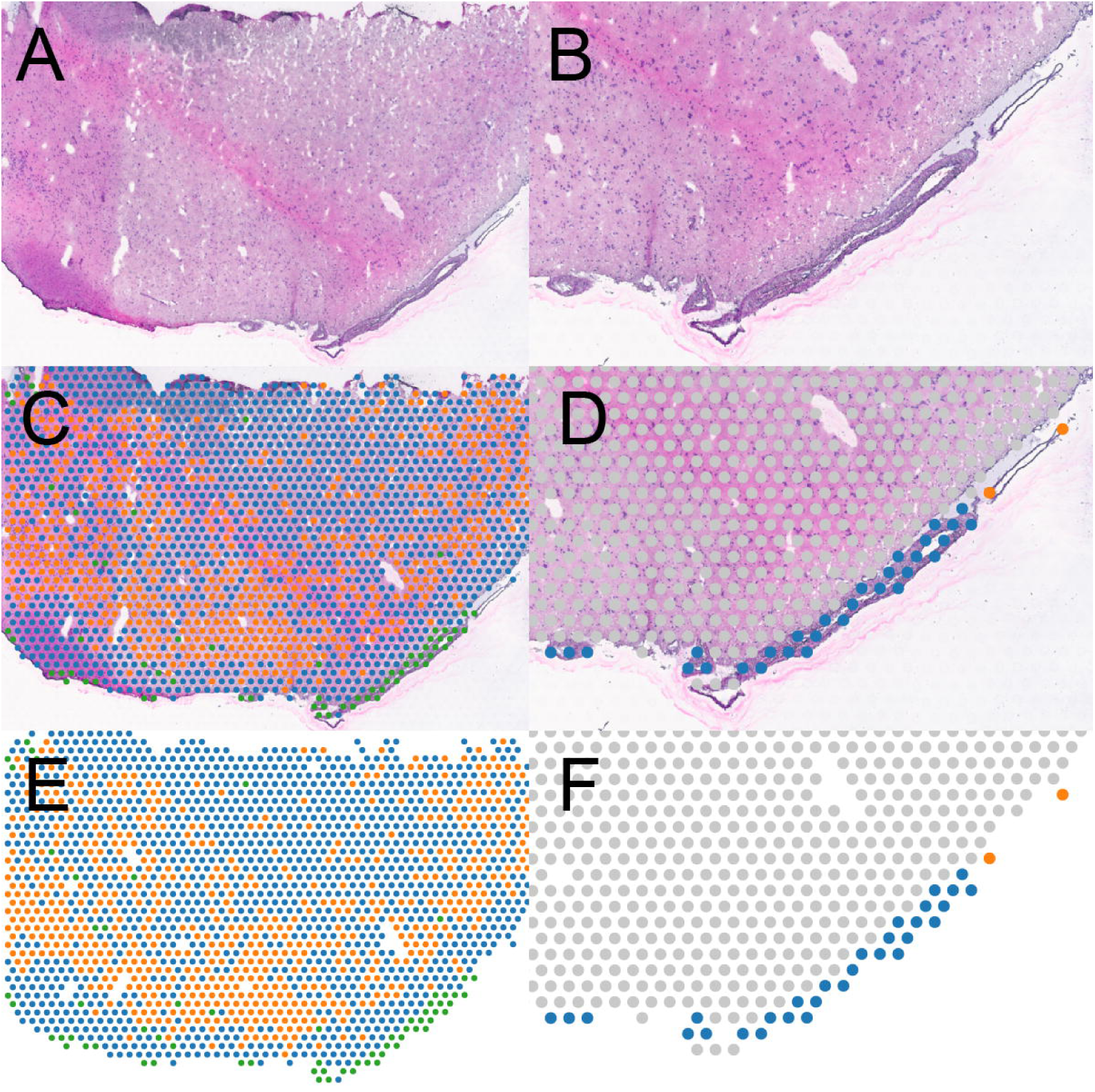
Loupe Browser display for SRT analysis of Block 2 with H&E (A,B), SRT dots for special foci (E,F) and fusion of both images (C,D). K-means clustering results are provided in the left pane (C,E) with green representing the cluster corresponding to vascular tissue and orange and blue corresponding to parenchymal tissues. Magnified view of dots remaining after tracing of vascular tissue based on histological appearance are provided in the right pane (D,F). Excluded dots are gray, while dots labeled artery are blue, and dots labeled vein are orange.

Differential expression analysis was performed, comparing dots labeled artery or vein based on visual appearance. Heatmaps for the top 50 differentially expressed genes from blocks 2 and 11B are provided in Figure 5 to demonstrate samples from the anterior and posterior circulations. Among the list of 50 differentially expressed genes for the blocks, the top four identified in both blocks are summarized in Figure 6, which provides details for the genes and their products, as well as expression differences and violin plots displaying these differences for each gene in each block. All four genes proved to refer to smooth muscle cells, which are present in arteries but not veins.

**Figure 5.**
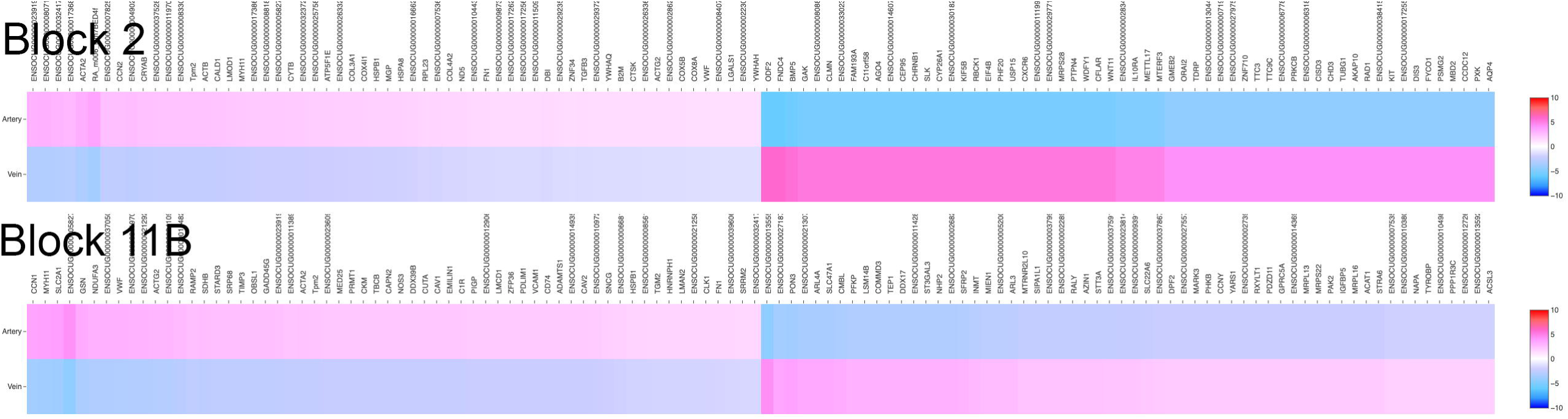
Heatmaps for the top 50 differentially expressed genes among the vessel-traced region, comparing arteries to veins for Blocks 2 and 11B. Differential expression between the groups are reflected in the color and shade, with increased expression indicated by red and blue for decreased expression. Gene IDs are listed above the heatmaps.

**Figure 6.**
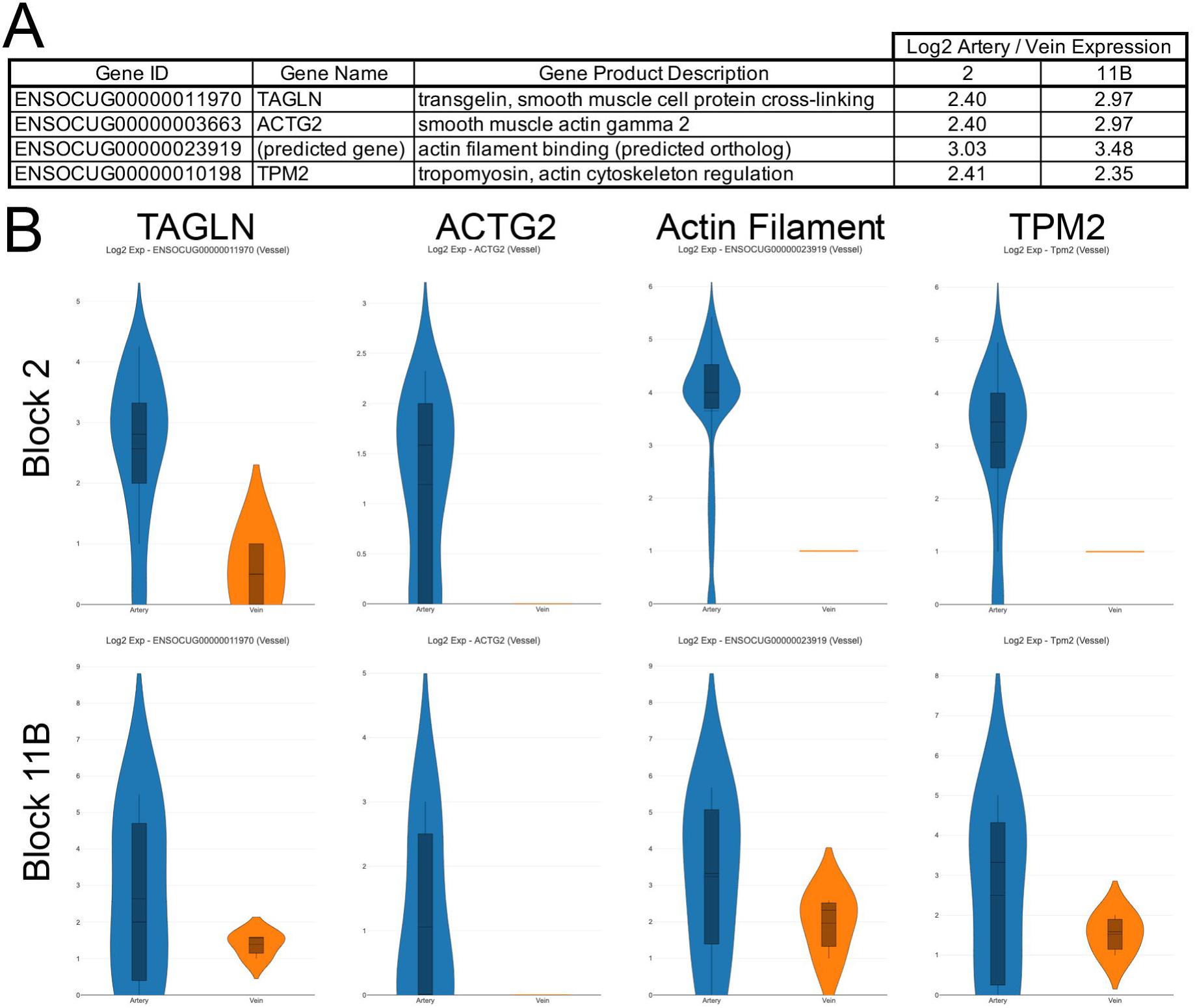
(A) Gene ID, gene name, and gene product description for four genes differentially expressed between arteries and veins with Log2 values of the differential expression ratio for each in the Blocks 2 and 11B. (B) violin plots for each of the 8 expression comparisons. Note each violin plot has unique scaling.

## Discussion

To develop more effective treatments for cerebrovascular disorders, it is imperative that we develop better understanding of vessel biology as it relates to the intracranial circulation specifically. To date, this has not occurred to a sufficient degree due to substantial obstacles to acquiring intracranial tissue. In such cases, animal models provide opportunities for understanding. There is a robust history of effective research performed using rabbits for investigation of cardiovascular diseases.(18–31)

The current report describes the amenability of rabbits to SRT methods for next generation evaluation of cerebrovascular disorders. To our knowledge, this represents the first report of application of the Visium SRT platform to either rabbits or the intracranial circulation. Data obtained from this model can be readily applied to human investigations given the amenability of both rabbits and humans to endovascular biopsy and performance of single-cell RNAseq on harvested cells.(26–30) SRT and single-cell RNAseq datasets are mutually reinforcing given the spatial data provided by the former and information obtained from intact cells obtained through a low-morbidity procedure in the latter.

The bioinformatics analysis performed on the samples in the present study yield face validity for the amenability of rabbit cerebrovascular tissues to SRT. Unsupervised machine-learning clustering analysis independently localized vascular tissue based on gene expression profiles without bias from anatomic considerations or other sources of bias that could be introduced by a human interpreter. Furthermore, utility for identification of tissue type, specifically distinguishing arteries from veins, was demonstrated by comparing differential expression traits between these vessel types. In cursory analysis of the most pronounced differentially expressed genes when comparing between the vessel types, the four most differentially expressed genes were related to smooth muscle cells, which are present in arteries but absent in veins. This finding augments the face validity of these methods for study of intracranial tissues like vessels.

While the present study describes the feasibility of SRT techniques for an animal model of cerebrovascular disorders, several limitations bear discussion. These results reflect preliminary investigation, and suboptimal tissue slice orientations limited the amount of vascular tissue present for analysis and the plane through which they were sliced. While all samples met quality control standards at each step, some samples demonstrated artifacts like fragmentation or curling. While all tissue is prone to such artifacts, further optimization of sample isolation and freezing may be possible to better limit such findings in rabbit brain samples. Finally, while providing gene expression data for thousands of locations across four samples, this still represents a dataset of limited size intended for proof-of-concept evaluation. As such, findings from cursory bioinformatics analysis of these data are not generalizable to make reach firm conclusions about vessel biology. Given the feasibility of these methods for analysis of rabbit models, further investigation is warranted to begin gathering data to obtain more impactful results.

## Conclusion

Animal models can play an integral role in acquiring better understanding of the vessel biology underlying the intracranial vasculature and the diseases it can develop. SRT using the Visium platform can be performed on samples from such animals to effectively isolate gene expression data enriched with spatial information. Further investigation utilizing these methods is warranted to further refine these potentially transformative techniques.

## Acknowledgements

This work was funded in part by The Joe Niekro Research Grant from the Joe Niekro Foundation and Society for Neurointerventional Surgery Foundation, American Heart Association Transformational Grant (19TPA34910194), and The Mark H. Huntsman Endowed Chair for Research in the Department of Radiology and Imaging Sciences at the University of Utah

## No conflicts of interest

All authors made a substantial contribution to the concept or design of the work, data acquisition, and analysis and interpretation of data. All authors drafted the article or revised it critically for important intellectual content and approved the version to be published. Each author participated sufficiently in the work to take public responsibility for appropriate portions of the content.

## Notes

Conflicting Interests: None.

### Competing Interest Statement

The authors have declared no competing interest.

